# Transcriptome-wide mapping of small ribosomal subunits elucidates scanning mechanisms of translation initiation in the mammalian brain

**DOI:** 10.1101/2024.11.07.622404

**Authors:** Preeti M Kute, Francois P Pauzin, Kornel Labun, Clive R Bramham, Eivind Valen

**Author notes:** shared first authors (Preeti M Kute, Francois P Pauzin, Kornel Labun).

## Abstract

Protein synthesis in neurons is highly compartmentalised and regulated, with key roles for translation initiation and elongation factors. The most widely used transcriptome-wide method for measuring translation, ribosome profiling, characterises the elongation phase of translation but does not provide insight into the initiation phase with scanning of the small ribosomal subunit (SSU). Here, we adapted and optimised ribosome complex profiling (RCP-seq) for brain tissue, capturing SSUs and analysis of translation initiation dynamics in mouse dentate gyrus and cerebral cortex. In both tissues, SSUs accumulate upstream of the start codon on synaptically localised RNAs and this ‘poised’ SSU configuration is associated with enhanced translational efficiency. Upstream open reading frames (uORFs) are extensively translated and associated with less SSU poising downstream, suggesting that uORFs may have a buffering effect on poised SSUs. Ribosome occupancy analysis suggests that neuron-specific transcripts recruit more ribosomes and are more translated than glia-specific transcripts. Furthermore, monosome-preferring neuronal mRNAs exhibit reduced scanning and elongation relative to polysome-preferring transcripts implying reduced recruitment of ribosomes. In sum, RCP-seq elucidates translation initiation dynamics in the mammalian brain and uncovers cell-type- and transcript-specific regulation.

## Introduction

Eukaryotic translation can be divided into four distinct stages: initiation, elongation, termination, and ribosome recycling. However, knowledge of translation dynamics across these stages is limited due to the lack of methods with nucleotide resolution in live systems (Hinnebusch 2014; Sokabe and Fraser 2019; Merrick and Pavitt 2018; Larsson, Tian, and Sonenberg 2013). Translation initiation, a rate-limiting process in protein synthesis, can be further divided into recruitment, scanning, and start codon recognition, which can all be subject to regulation. During recruitment, the assembly of the preinitiation complex (PIC) containing the small ribosomal subunit (SSU/40S) with methionyl-initiator tRNA, and subsequent recruitment to the mRNA 5’ cap, is regulated by eukaryotic initiation factors (eIFs) (Hinnebusch 2014). Following recruitment, the PIC scans the 5’ leader for a start codon, after which the large ribosomal subunit (LSU/60S) joins the SSU to form an elongation-competent ribosome (80S).

Translation in neurons is highly compartmentalised and regulated due to their complex polar structure and function. A prominent example is the local translation in the synaptic compartment remote from the cell body, important for synapse development, maintenance, and plasticity (Holt, Martin, and Schuman 2019; Bramham and Wells 2007; Sun and Schuman 2023). Both translation initiation and elongation in the nervous system are modulated by changes in translation factor activity in response to changes in neural activity and neurotransmitter receptor activation. Converging evidence highlights the role of translation initiation factors (eIF2α, eIF4E) as regulators of protein synthesis in synaptic plasticity and diverse functions such as circadian rhythms, social behaviour, memory, and cognitive flexibility (Patil et al. 2023; Amorim et al. 2018; Cao et al. 2015; Sharma et al. 2020; Moon, Sonenberg, and Parker 2018; Oliveira and Klann 2022). While these studies demonstrate the role of initiation factors in PIC recruitment and 5’ leader scanning, they do not resolve the dynamics of SSU scanning and start codon recognition.

Analysing 80S ribosome-bound mRNAs through ribosome profiling and related techniques such as TRAP-seq has provided insight into many questions related to translation control in the mammalian brain (Thomson et al. 2017; Hien et al. 2020; Amorim et al. 2018). However, ribosome profiling characterises only the elongation step of translation such that the dynamics and regulation of translation initiation in the mammalian nervous system have remained elusive. Translation complex profiling (TCP-seq) was the first method to characterise the occupancy of SSUs across the transcriptome (Archer et al. 2016). TCP-seq, like Ribo-seq, involves digesting the lysates with RNase I and separating 40S and 80S fractions based on the polysome profile obtained after sucrose gradient density centrifugation. RNA from these fractions is then sequenced and analysed. In the original TCP-seq study only those mRNAs bound by at least an elongating (80S) ribosome were used to explore translation initiation in yeast and a modified technique, ribosome complex profiling (RCP-seq) was developed to capture all mRNAs, including those bound only by a 40S with no elongating (80S) ribosomes, to explore initiation in the early development of zebrafish (Giess et al. 2020). In the mammalian brain, however, transcriptome-wide characterization of translation initiation has not been attempted.

Here we developed a UV light crosslinking method capable of producing sufficient input material for RCP-seq for brain tissues from mice. This approach generated libraries of similar quality to previous studies (Archer et al. 2016; Giess et al. 2020) and allowed a detailed characterization of translation initiation and elongation in the nervous system for the first time. Using the dentate gyrus region of the hippocampus and the cerebral cortex from mice, we show regulation of translation at the initiation stage. First, SSUs accumulate upstream of the start codon in a poised state on synaptically localised mRNAs, indicating a regulatory step during the transition to the elongation step. Second, our data reveals regulation during scanning with a role for uORF-mediated translational repression of the downstream CDS. Third, we show that neuron-enriched transcripts, relative to their RNA abundance, have a higher recruitment of ribosomes leading to enrichment of both scanning and elongating ribosomes as compared to glia-enriched transcripts. Finally, we show that neuronal transcripts preferring monosomal translation have reduced recruitment of ribosomes, causing their reduced translation efficiency.

## Results

### Mapping the SSU and 80S ribosomal complexes in the dentate gyrus (DG) and cerebral cortex of the adult mouse brain

Previous methods for capturing scanning SSUs have used formaldehyde and chemical crosslinking for the SSU fixation to obtain footprints on the mRNAs (Archer et al. 2016; Bohlen et al. 2020; Giess et al. 2020; Wagner et al. 2020). In mouse brain tissue, the use of 0.1% or 0.05% formaldehyde resulted in less crosslinking for polysomes as compared to UV-crosslinked lysates (**Supplementary Figure 1a**). Thus, we opted for UV-crosslinking, which has been previously used for studying RNA binding protein (RBP) interactions with mRNAs (Hafner et al. 2021) and for polysome profiling in the brain tissue (Darnell et al. 2011). The steps to efficiently crosslink the SSU and the 80S on RNAs from the mouse brain tissue are outlined in **Figure 1a** and detailed in the methods section. Briefly, the pre-cleared lysate was exposed to UV irradiation and digested with RNase I to obtain footprints (**Supplementary Figure 1b**). Since the SSU peak was undetectable in the polysome profiles, RNA was isolated from DG polysome profiling fractions and run on a Bioanalyzer to detect the presence or absence of the 28S rRNA to differentiate SSU from the 80S fractions (**Supplementary Figures 1c and 1d**). The fractions corresponding to the SSU and 80S were collected post-digestion and libraries were prepared and sequenced to 20-130 million reads per library (**Supplementary Table 1**). Total RNA libraries were prepared from matched undigested samples to estimate RNA abundance (**Supplementary Table 1**).

**Figure 1:**
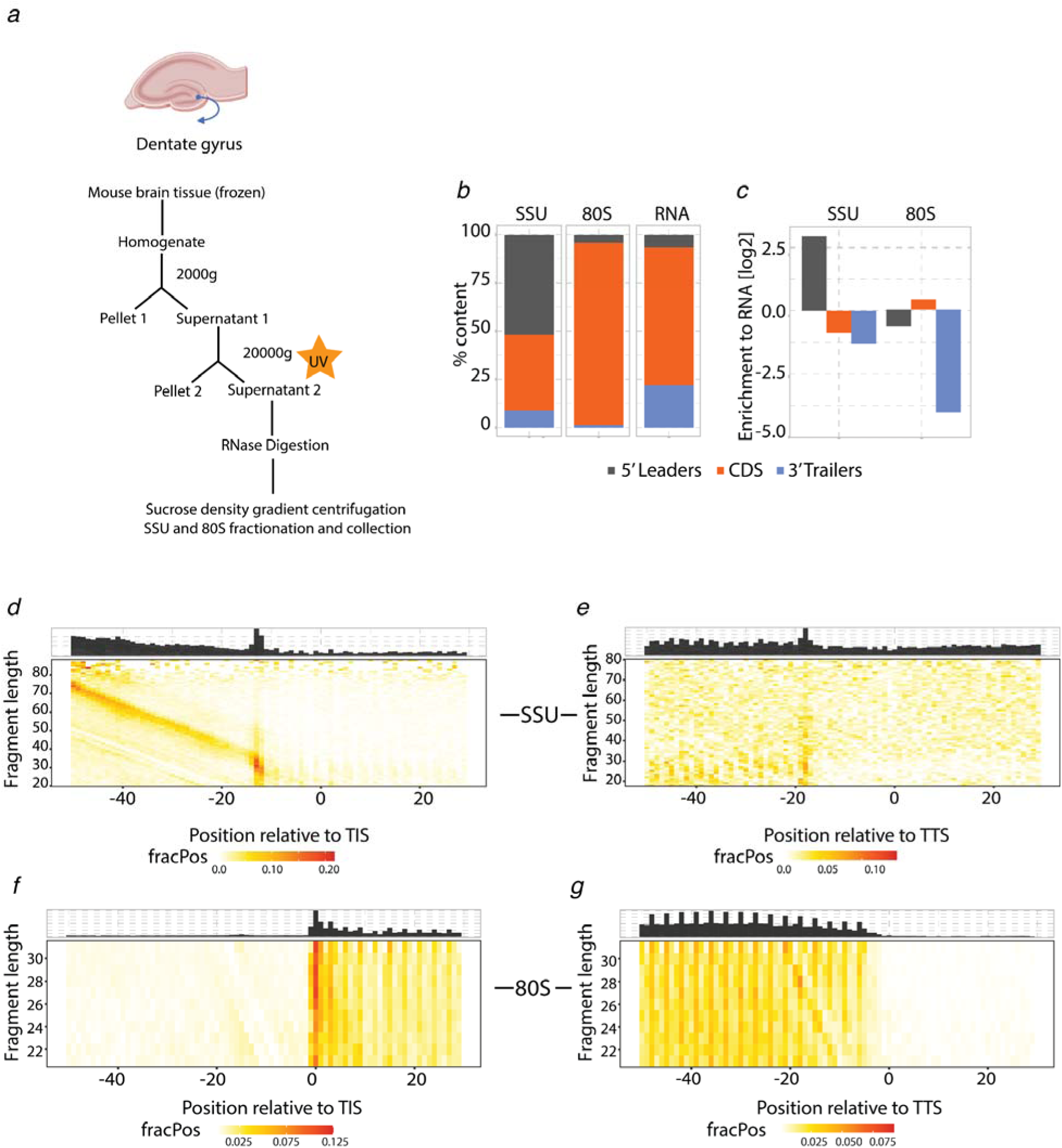
Ribosome complex profiling (RCP-seq) captures small ribosomal subunits (SSU) and elongating ribosomes (80S) from the mouse DG tissue. **a**: Steps for ribosomal complex profiling using UV crosslinking for the fixation of the SSU and 80S in the DG. **b** and **c**: Content plots displaying the mapping of SSU, 80S, and total mRNA counts to the 5’ leaders, CDS, and 3’ trailers of the transcripts (n = 3). **b)** shows the % count for each library and **c)** displays the normalised counts to the total RNA. **d**-**e**: Footprint length distribution for the 5’ end of SSU fragments in the DG for **d)** translation initiation site (TIS) and **e)** translation termination site (TTS). FracPos indicates the fraction of counts per position per gene. **f**-**g**: Footprint length distribution for the 5’ end of 80S fragments in the DG for **f)** translation initiation site (TIS) and **g)** translation termination site (TTS). FracPos indicates the fraction of counts per position per gene.

The library quality was comparable to previous studies (Archer et al. 2016; Giess et al. 2020) where contaminants were largely from rRNAs (**Supplementary Figures 1e**). Of the SSU reads mapped to mRNAs, 52% mapped to the 5’ leader (**Figure 1b**) while 94% of the 80S reads mapped to the coding sequence (CDS) (**Figure 1b**). Accordingly, when normalised to the expected fraction of reads from a given region, estimated from the total mRNA library, the SSU libraries and 80S libraries were enriched as expected in the 5’ leader and CDS respectively (**Figure 1c**). Footprint length distribution for the 80S libraries was enriched within the expected range: 26-30 nt (Ingolia et al. 2009) and showed longer reads up to 60 nt, implying either the presence of SSUs with initiation factors as discussed in a previous TCP-seq study (Wagner et al. 2020) or ribosomal interactions with other RNAs such as lncRNAs (Ruiz-Orera and Albà 2019). Similarly, SSU footprints had a broader distribution with many longer fragments, 20-75 nt in these libraries (**Supplementary Figure 1g**) as previously reported (Giess et al. 2020; Archer et al. 2016). Metagene heatmap of SSU footprints over the translation initiation site (TIS) for the DG showed a range of fragment lengths (20-75 nt) forming a diagonal preceding the TIS (**Figure 1d**). This has been attributed to longer pre-initiation complex conformations due to the presence of initiation factors (Bohlen et al. 2020; Archer et al. 2016). Some SSU footprints are also present internally in the CDS, seen upstream of TTS (**Figure 1e**), potentially representing contamination from dissociated 60S subunits during sample preparation or instances of “leaky scanning” where the PIC scans through the start codon and into the CDS. For the 80S ribosomes (**Figures 1f and 1g**), the footprints are enriched at the TIS and show the expected 3-nucleotide periodicity throughout the CDS. Although the reads mapping specifically to either 5’ leaders or CDS were low for SSU and 80S libraries (**Supplementary Figures 1f**) replicates were highly correlated (**Supplementary Figures 1h-1j**), and clustered separately as SSU, 80S, and RNA (**Supplementary Figure 1k**) with PCA, demonstrating that the protocol is robust and reproducible.

To assess the transferability of the technique to another brain region, we applied RCP-seq to cerebral cortex tissue (**Supplementary Figures 2a-d**). The library quality was comparable to the DG, with 37% of the SSU reads and 94% of the 80S reads of mRNA reads mapping to the 5’ leaders and CDS respectively (**Figure 2a** **and Supplementary Figure 2e**) with a strong enrichment relative to the expected distribution based on RNA (**Figure 2b**). Footprint length distributions were within the expected range of 20-75 nt but more abundant between 25-30 nt for SSU and 28-32 nt for 80S, with some reads going up to 60 nt (**Supplementary Figure 2f**). As compared to DG, cortical SSU showed less enrichment of longer footprint lengths (>40 nt), which could be attributed to either a technical difference or biological difference between the two tissue types. As expected, the metagene heatmap of SSU footprints showed a diagonal of fragments (20-75 nt) preceding the TIS (**Figure 2c**), which is attributed to a longer pre-initiation complex conformation due to the presence of initiation factors (Bohlen et al. 2020; Archer et al. 2016). Surprisingly, we observed an additional diagonal pattern from ∼10nt longer reads indicating a second SSU PIC conformation at the TIS (**Figures 2c**). This suggests that additional factors such as eIF3B, eIF4G1 could be involved in the formation of this SSU configuration as discussed previously (Bohlen et al. 2020). Some SSU footprints are also present internally in the CDS, as seen in DG (Figure 2d). For the 80S ribosomes, the footprints are enriched at the TIS and TTS and show the expected 3-nucleotide periodicity throughout the CDS (**Figures 2e and 2f**).

**Figure 2:**
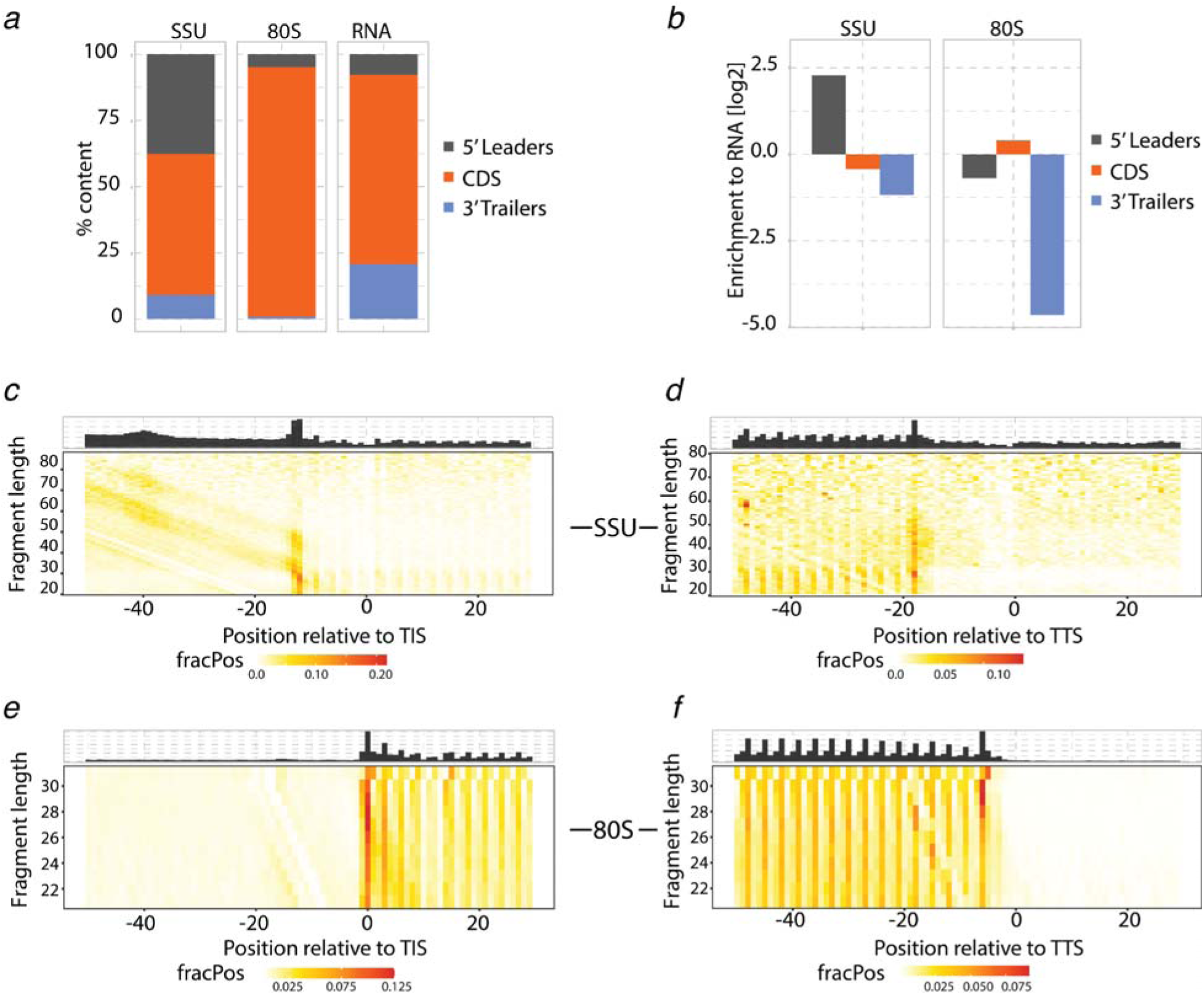
Ribosome complex profiling (RCP-seq) captures small ribosomal subunits (SSU) and elongating ribosomes (80S) from the mouse cortical tissue. **a** and **b**: Content plots displaying the mapping of SSU, 80S, and total mRNA counts to the 5’ leaders, CDS, and 3’ trailers of the transcripts (n = 3). **a** shows the % count for each library and **b** displays the normalised counts to the total RNA. **c and d**: Footprint length distribution for the 5’ end of SSU fragments in the cortex for **c)** translation initiation site (TIS) and **d)** translation termination site (TTS). FracPos indicates the fraction of counts per position per gene. **e and f**: Footprint length distribution for the 5’ end of 80S fragments in the cortex for **e)** translation initiation site (TIS) and **f)** translation termination site (TTS). FracPos indicates the fraction of counts per position per gene.

Taken together, our UV-crosslinking method successfully captures the SSU and 80S ribosomal complexes transcriptome-wide in the adult mouse DG region of the hippocampus and in the cerebral cortex.

### Poised SSUs upstream of TIS in selected genes

Further analysis of the start-codon-associated SSU footprints showed two distinct populations of SSUs with different 5’ ends but overlapping 3’ ends (**Figure 3a**, **Supplementary Figure 3a and 3b**). The majority of the SSU population had a 5’ end at -12 with 3’ ends at +11 to +25 similar to those observed for SSUs from HEK293T (Wagner et al. 2020). This SSU position implies a closed conformation of SSU following AUG recognition (Archer et al. 2016) (**Figure 3b**, top row). The longer SSU footprints have previously been explained as pre-initiation complexes associated with initiation factors (IFs) that extend the protection of the RNA (Archer et al. 2016; Giess et al. 2020; Bohlen et al. 2020; Wagner et al. 2020). These longer SSU footprints with extended 5’ ends at -46 to -36 nt (Supplementary Figures 3a and 3b, blue lines) share 3’ ends with the shorter conformation that lack IFs and does not continue scanning further downstream. These are therefore “poised” to initiate elongation (possibly in a post-AUG-recognition conformation). Besides SSUs with IFs (**Figure 3b**, middle row) the length of “poised” is also consistent with two queued SSUs (**Figure 3b**, bottom row). Previous TCP-seq studies have speculated that these longer SSU fragments can arise due to queuing of multiple SSUs near the start codon (Archer et al. 2016; Wagner et al. 2020). Such queued SSUs were also demonstrated to occur *in vitro* (Shirokikh et al. 2019) and posited in various studies (summarised in Supplementary Table 2). To rule out that UV-crosslinking specifically enriches these SSU conformations in the brain tissues, we performed formaldehyde crosslinking in HEK293T cells following the Wagner et al protocol (Wagner et al. 2020) and compared it to UV crosslinking. In formaldehyde crosslinked samples, we observed a similar enrichment of SSUs with 5’ ends around -50nt relative to the TIS (**Supplementary Figures 3d**), while absence of crosslinking gave no such enrichment of SSU footprints (**Supplementary Figures 3c**). UV crosslinking before and after lysis also showed a similar but lesser enrichment of SSUs with 5’ ends around -50nt and relative to the TIS (**Supplementary Figures 3e-3f**), indicating that similar SSU conformations are observed both in HEK293T cells and brain tissues, irrespective of the crosslinking method.

**Figure 3:**
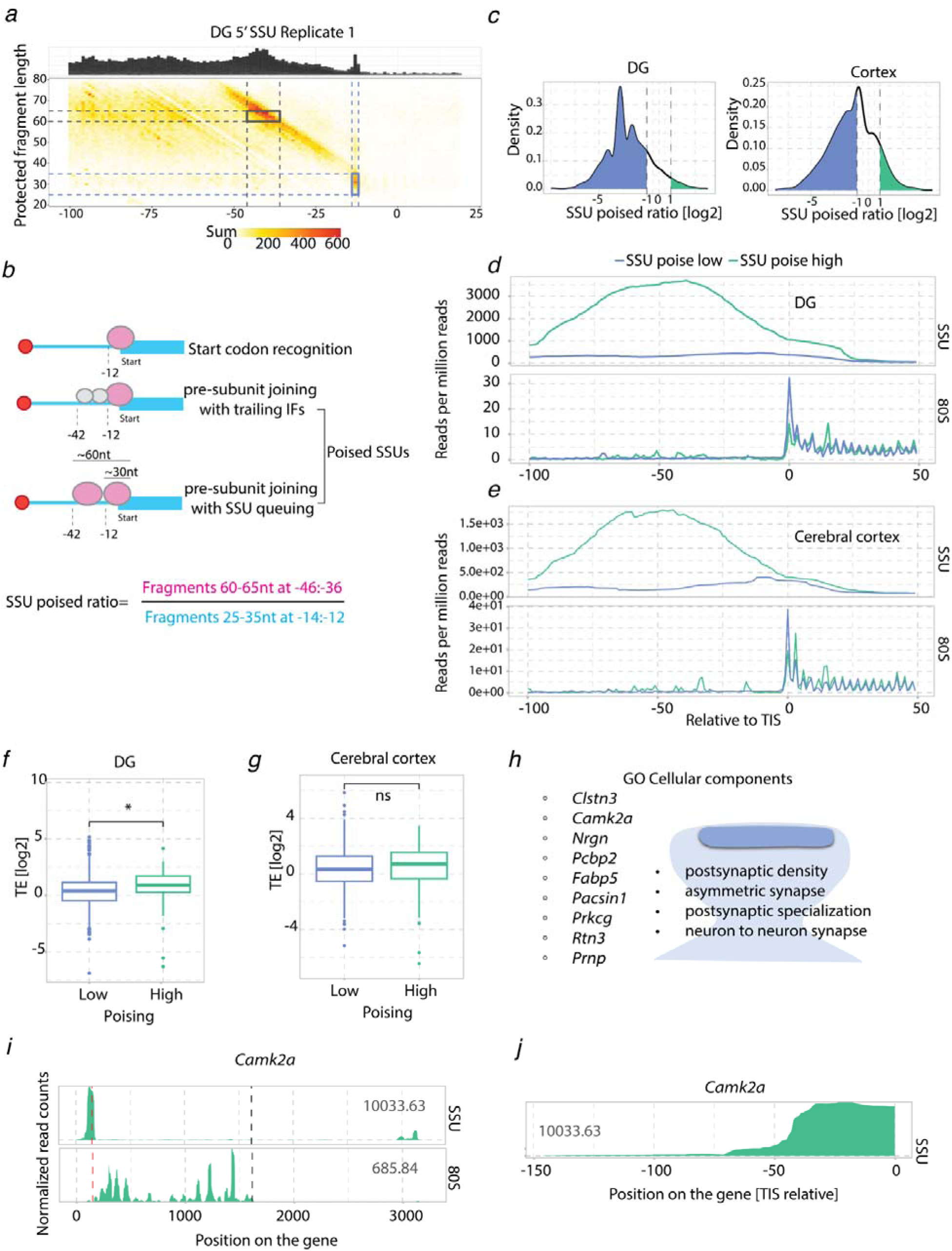
SSU build-up observed in the leaders of transcripts in the DG and the cerebral cortex. **a**: Footprint length distribution for the 5’ end of SSU fragments in the DG for the TIS, highlighting initiating SSUs (at -12nt) and poised SSUs (-46: −42nt). **b**: Schematic of different conformations of SSUs near the TIS of transcripts. Formulae for calculating SSU poised ratio. **c**: Distribution plot for SSU poised ratio (log2) for DG and cortical transcripts. Dotted lines indicate SSU poised ratio cut-off of - 1 and +1. Coloured regions indicate transcripts with lower (blue) or higher (green) poised ratios from the cut-off lines. **d**: Metagene coverage for transcripts with SSU poised ratio below (blue) or higher (green) than the cut-offs in the DG relative to the TIS. **e:** Metagene coverage for transcripts with SSU poised ratio below (blue) or higher (green) than the cut-offs in the cerebral cortex relative to the TIS **f**: Log_2_ of Translation Efficiency (TE) for genes with high or low poised ratios (FPKM based, t-test statistics) for DG transcripts. **g**: Log_2_ of Translation Efficiency (TE) for genes with high or low poised ratios (FPKM based, t-test statistics) for cortical transcripts. **h**: GO enrichment categories for genes with SSU poising ratio more than 2 and the list of genes enriched in the categories in both DG and cortex. **i**: Single gene profile for SSU and 80S, without intronic regions, and plotted for the longest isoform of *Camk2*α from DG. Numbers indicate FPKM values. Dotted lines indicate TIS (red) and TTS (black). **j**: Single gene profile for SSU within the 5’ leader region of *Camk2*α from DG. Numbers indicate FPKM values.

We next asked where and to what extent poised SSUs occur in the transcriptome. As poised SSUs overlap the start codon, they cannot coexist on the same transcript with the short SSU conformations at - 12 nt that initiate elongation. The transition between these two conformations could therefore indicate a regulatory step before or during start codon recognition concomitant with a change in ribosomal conformational state. We, therefore, defined an “SSU poised ratio” as the ratio between the two mutually exclusive configurations of poised SSUs and initiating SSUs (boundaries in **Figures 3a and 3b**) and calculated this metric for 2725 transcripts that had sufficient coverage of SSU reads, from the cortical and DG tissues. This revealed that most transcripts have more initiating SSUs than poised SSUs, but that a subset of transcripts have substantially higher poising (**Figure 3c** below *vs*. above zero). The length of the 5’ UTRs can influence the density of SSUs <u>(Archer et al. 2016)</u> that could cause SSU accumulation on shorter 5’UTRs. Interestingly, the length of 5’ UTRs was not a contributing factor to the extent of SSU poising (**Supplementary Figure 4a)**. We selected the strongest SSU-poised transcripts as those having a two-fold enrichment in poised *vs*. initiating SSUs (76 genes for DG and 198 for cerebral cortex) and the weakest as two-fold depletion (1536 genes for DG and 1041 for cerebral cortex). This revealed that poised genes have substantially more scanning activity with a higher occupancy of SSUs across the 5’ leader, but a weaker presence of 80S ribosomes at the TIS (**Figures 3d and 3e**). This could indicate increased recruitment of PIC to the RNA, potentially combined with either a slower joining of 60S or a faster translocation of 80S away from the TIS. We calculated the translation efficiency (TE, see Methods) metric to assess the presence of 80S on these transcripts and observed higher TE for transcripts with a high poising ratio as compared to those with a low poising ratio for DG but not for the cerebral cortex (**Figures 3f and 3g**).

To probe the biological relevance of poised SSUs, we analysed the 42 genes poised in both DG (76 genes) and cortex (198 genes) tissues for overrepresented functional GO categories against a restrictive background of other genes with sufficient reads, in total 2725 genes (1917 DG, 1969 cortex) (**Supplementary Table 3**). This revealed four enriched categories (**Figure 3h**) all belonging to the synaptic compartment of the neuron. A prominent gene is *Camk2*α (**Figures 3i and 3j**) where the SSU occupancy plot showed a strong enrichment of SSUs near and up to 60 nt upstream of the TIS (**Figure 3j**). Other examples include *Clstn3*, *Pcbp2*, *Pacsin1*, *Rtn3*, *Prnp*, and *Prkcg* (**Supplementary Figure 3g-3l**). Together this shows that poised SSUs are enriched in a subset of functionally related genes.

### uORF-mediated regulation in the DG tissue

Upstream open reading frames (uORFs) are regions in the 5’ leaders of mRNAs with a start codon and a downstream in-frame stop codon, which could potentially encode a peptide. While many of these uORFs are translated, it is believed that most of these do not produce functional peptides but rather contribute to regulating translation of the downstream CDS (Dever, Ivanov, and Hinnebusch 2023; Kute et al. 2021). A key regulatory feature of uORFs is that some ribosomes detach after translating the uORF and are therefore unable to initiate at the protein-coding TIS (**Figure 4a**), leading to translation downregulation of the CDS. To explore the role of uORFs in the translation regulation of downstream CDSs in the DG, we defined uORFs as ORFs with 80S read coverage and having an AUG as the start codon. For those 5’ leaders containing multiple eligible uORFs we selected the one with the highest read coverage (totaling 1062, see methods). These were termed “active uORFs” (**Supplementary Table 4**). To check the effect of the translation of active uORFs on downstream CDS, we looked at the effect of the presence or absence of uORFs on the translation of the transcripts. This showed a modest but significant inhibitory effect of having actively translated uORFs (**Figure 4b**). To further explore the role of uORFs in regulation of CDS translation we calculated the scanning efficiency (SE) (**Figure 4c**) which quantifies the abundance of SSU occupancy in the leaders normalised to the RNA levels. SEs were comparable between transcripts harbouring uORFs and those without, implying that uORFs do not globally have a significant effect on the overall occupancy of SSUs over the leaders (**Figure 4d**). On the other hand, uORFs can affect the SSU traversal over the 5’ leaders either through slowing them down by the relatively slower process of elongating a uORF or by dissociating a proportion of them from the RNA after uORF translation. To check whether uORFs had any impact on the number of SSUs reaching the TIS, we looked at poised SSUs at the position -40 nt and length 60 nt for genes with and without uORFs. The poised SSUs reads were normalised to RNA to accommodate any differences at the RNA level. This analysis revealed that the poised SSUs in those genes containing an active uORF were significantly lower (**Figures 4e and 4f**) suggesting that uORFs may act to slow down or disassociate SSUs on their way to the CDS. We next asked whether the presence of uORFs leads to SSU poising at the TIS of the uORF. However, we did not observe this to the same extent as for protein-coding TISs (**Supplementary Figure 4b**). This may be because uORF initiation is typically more stochastic than CDS initiation and that leaky scanning (i.e. not initiating at the uORF TIS) may lead to SSUs spread between, potentially, multiple uORFs and the CDS.

**Figure 4:**
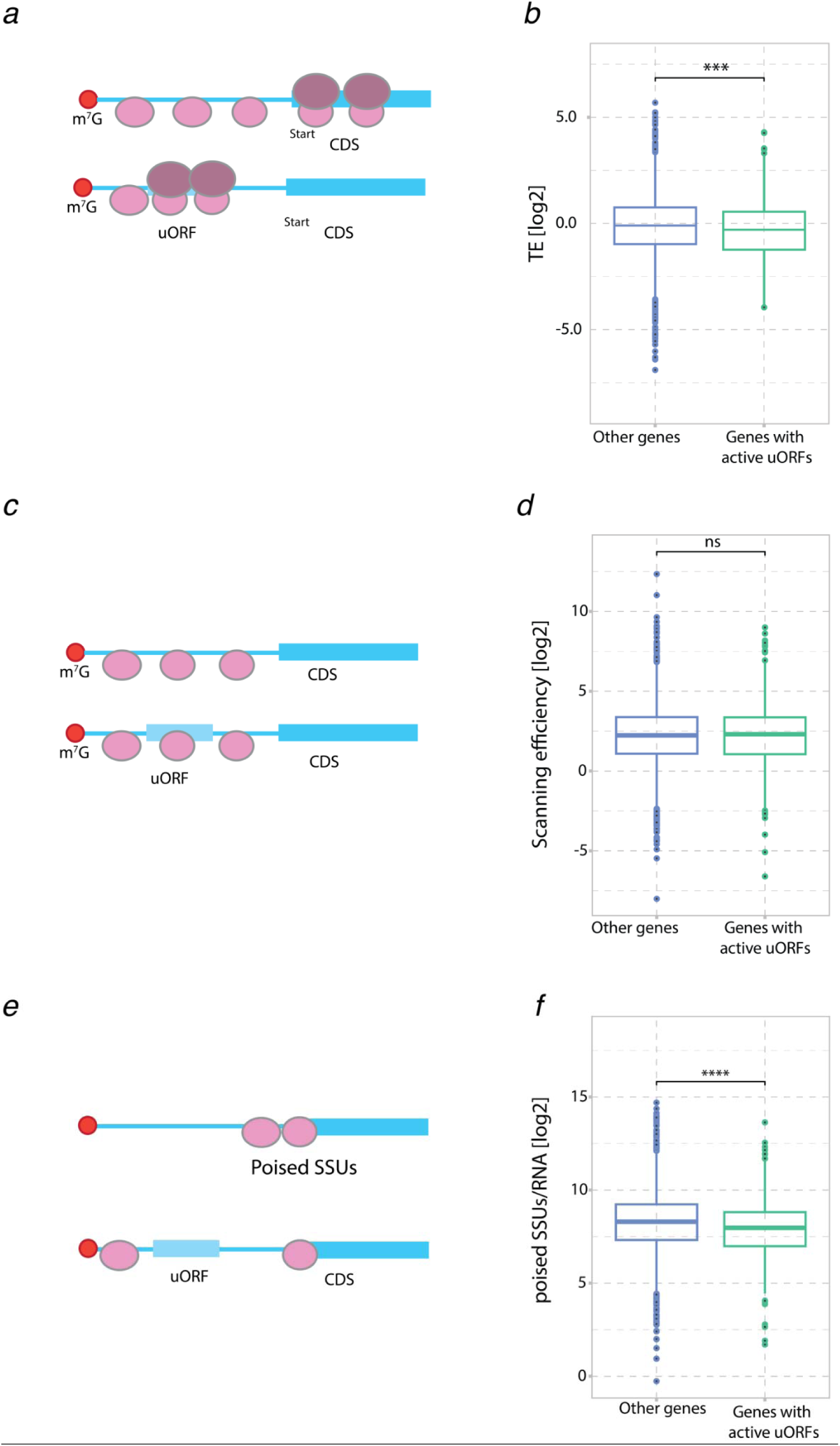
uORF expression in the DG. **a**: Illustration explaining the role of uORFs on the translation of downstream CDS. **b**: Comparison of CDS translation efficiency (TE) in relation to the presence or absence of active uORFs in DG (FPKM based, t-test statistics). **c**: Illustration explaining the role of uORFs on the scanning efficiency (SE) of downstream CDS **d**: Comparison of the SE in relation to the presence or absence of active uORFs in DG (FPKM based, t-test statistics). **e**: Illustration explaining the role of uORFs on the SSU poising over 5’ leaders **f:** Comparison of the poised SSUs in relation to the presence or absence of active uORFs in DG (FPKM based, t-test statistics).

### The translational landscape of neuronal and glial genes

The mammalian brain consists of both neuronal and non-neuronal cells such as astrocytes, oligodendrocytes, microglia, and blood vessel cells. To elucidate the translational landscape of cell-type specific transcripts, we filtered our dataset according to previously defined neuronal and glial-enriched transcripts (Glock et al. 2021). Neuronally-enriched transcripts show more coverage of SSU, 80S and RNA as compared to glial-enriched transcripts which may be reflective of the cell-type abundances in the tissue (**Figure 5a**). Taking into account these differences by normalising by RNA abundance, neuronal-enriched transcripts showed modest but significantly higher TE and SE values as compared to glial-enriched transcripts (**Figures 5b and 5c**). The gene profiles for neuronal transcripts *C1ql2* and *Prox1* (**Figures 5d and 5e**) exemplifies the differences in SSU and 80S occupancy compared to glial-enriched transcripts *Tgfb2* and *Fth1* (**Figures 5f and 5g**). Taken together, the neuronal-enriched transcripts are scanned and translated more efficiently than glial-enriched transcripts. This suggests a cell-type-specific difference, with enhanced translation of neuronal transcripts relative to glial transcripts in the DG.

**Figure 5:**
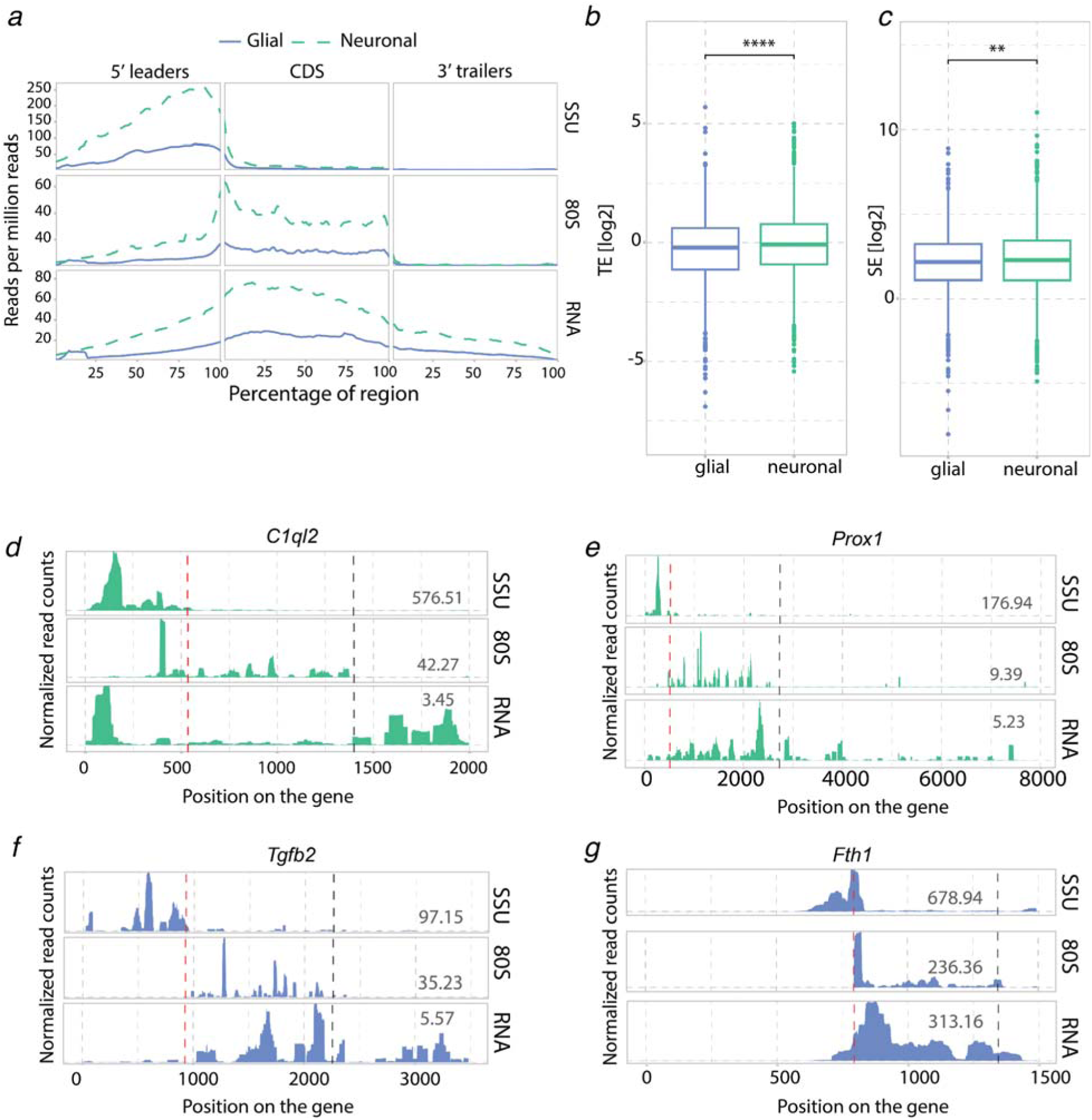
Transcriptional and translational profiles for neuronal and glial-specific genes in the DG of mouse hippocampus. **a**: Metagene coverage for the SSU, the 80S and total RNA between neuronal- and glial-enriched transcripts. **b and c**: Translation efficiency and scanning efficiency for glial and neuronal-specific transcripts ((FPKM based, t-test statistics) **d-g**: Single gene profiles for SSU, 80S and total RNA coverage, without intronic regions, and plotted for the longest isoform. **d**) *C1ql2* (Complement Component 1, Q Subcomponent-Like 2) , **e**) *Prox1* (Prospero homeobox protein 1), **f**) *Tgfb2* (Transforming growth factor beta 2), **g**) *Fth1*(Ferritin heavy chain 1). Numbers indicate FPKM values. Dotted lines indicate TIS (red) and TTS (black).

### Scanning and elongation in neuropil-enriched monosome- *versus* polysome-preferring transcripts

Since translation in neurons is compartmentalised, occurring even in distal dendrites, we next studied transcripts that are differentially localised and translated between the soma and the synaptic neuropil. In a previous study based on ribosome profiling, more than 800 transcripts were found to be predominantly translated in the neuropil (neuronal dendrites and axons) of microdissected rat hippocampal tissue slices (Glock et al. 2021). Surprisingly, a recent report based on polysomal profiling of microdissected hippocampal CA1 tissue showed that most of the neuropil translation is mediated by monosomes, with only one actively translating ribosome per transcript (Biever et al. 2020) (**Figure 6a**). The authors also noted that monosome-preferring transcripts had lower translation initiation and elongation rates. Here we used RCP-seq to directly assess initiation and elongation in neuronal-specific transcripts preferring monosomal or polysomal translation (Glock et al. 2021; Biever et al. 2020). Notably, for both soma and neuropil compartments, monosome-preferring transcripts showed lower SSU, 80S, and RNA abundance as compared to those that preferred polysomal translation (**Figures 6b and 6c**). Interestingly, while monosome-preferring transcripts showed lower TE than polysome-preferring transcripts from the neuropil compartment (**Figure 6d**), they displayed similar levels of occupancy from scanning ribosomes (**Figure 5e**) coupled with a lower level of initiation (**Figure 6f**).

**Figure 6:**
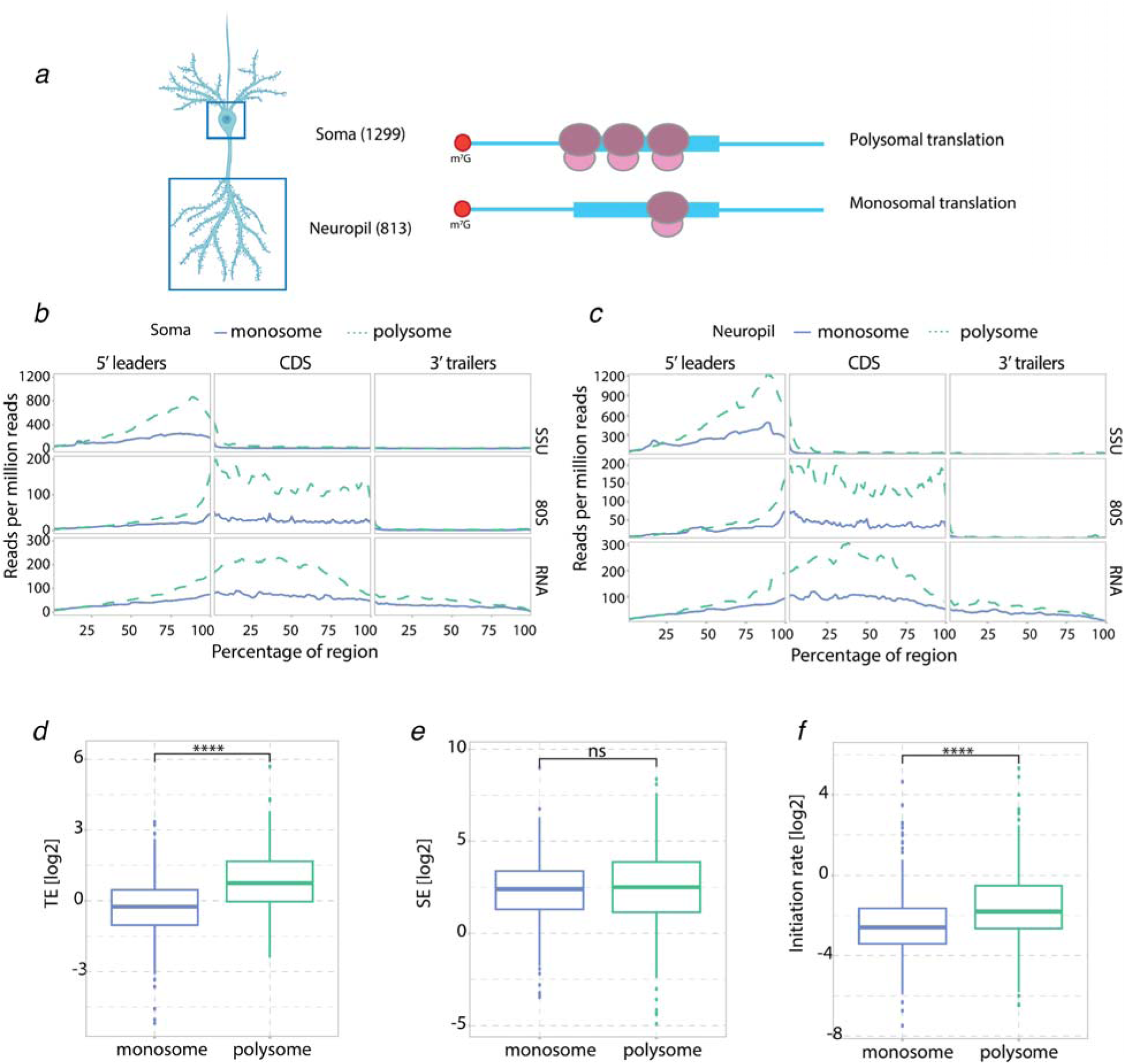
Translation landscape of neuropil and soma-enriched transcripts. **a**: Representation of neuronal compartments (soma and neuropil) with polysomal translation (multiple ribosomes elongatin simultaneously) and of monosomal translation (only one ribosome elongating on the transcript). Numbers in parentheses indicate the number of genes used for analysis **b**: Metagene coverage for transcripts enriched in the soma and preferring monosomal or polysomal translation. **c**: Metagene coverage for transcripts enriched in the neuropil and preferring monosomal or polysomal translation. **d**: Log_2_ TE (FPKM based, t-test statistics) for neuropil-enriched transcripts between the transcripts preferring monosomal or polysomal translation in the DG. **e**: Log_2_ SE (FPKM based, t-test statistics) for neuropil-enriched transcripts between the transcripts preferring monosomal or polysomal translation in the DG. **f**: Log_2_ Initiation rate (FPKM based, t-test statistics) for neuropil-enriched transcripts between the transcripts preferring monosomal or polysomal translation in the DG.

## Discussion

### Optimising RCP-seq for brain tissue

This study presents the first map of translation initiation dynamics in tandem with elongation in the mammalian brain. This was accomplished by adapting and optimising several steps of our previously published RCP-seq protocol for developing zebrafish embryos (Giess et al. 2020). While previous studies mapping SSUs to the mRNAs have used PFA and chemicals such as dithiobis (succinimidyl propionate, DSP) (Wagner et al. 2022; Giess et al. 2020), the use of low (0.05%) PFA concentrations in the brain tissues gave poor yield of polysomes, possibly due to insufficient lysis of the tissue. To increase the amount of SSUs and to avoid the use of glycine to neutralise formaldehyde, we crosslinked ribosomal complex proteins to RNA using UV irradiation, as used previously to map RBP binding sites and for polysome profiling (Hafner et al. 2021; Darnell et al. 2011). A possible drawback of UV crosslinking of lysates could be the loss of SSUs during the lysis step. However, in our data, we do not observe any indication of loss and see a distribution of SSUs over the leaders similar to previous studies (Archer et al. 2016; Giess et al. 2020; Wagner et al. 2022). UV irradiation is also known to cause ribosome stress response (M. S. Iordanov et al. 1998; Mihail S. Iordanov et al. 2002), but this study exposes the lysates to UV and thereby bypasses the harmful effects of UV irradiation on tissue. Together these steps enabled us to successfully capture the transcriptome-wide occupancy of both scanning and elongating ribosome complexes from the dentate gyrus and the cerebral cortex region of the mouse brain.

Our libraries of ribosomal occupancy from brain tissues were consistent with previously published datasets in terms of transcriptomic mapping, footprint length distributions, and 80S periodicity (Archer et al. 2016; Bohlen et al. 2020; Giess et al. 2020). Accordingly, we observed the diagonal preceding the TIS in the SSU libraries, which has been observed in RCP-seq and TCP-seq studies (Wagner et al. 2022; Giess et al. 2020). Interestingly, we observed a second diagonal in the cerebral cortex, indicating an approximately 10nt longer conformation present at the TIS. While sequencing data alone is insufficient to determine the factors present in this conformation, the read positions and lengths are indicative of the presence of one or more additional initiation factors downstream of the pre-initiation complex such as helicase eIF4A and its cofactor eIF4B as shown by recent structural evidence (Brito Querido et al. 2024).

### Role of uORFs and SSU-poising in neuronal translation

A major finding of this study was the characterization of a distinct poised configuration of SSUs enriched at the TISs of functionally related transcripts. Previous studies from yeast, developing zebrafish embryos and HEK293T cells have shown similar SSU patterns (Archer et al. 2016; Giess et al. 2020; Wagner et al. 2020) but not in Hela cell line (Bohlen et al. 2020). This could imply that poising is a species or context-specific phenomenon. We introduced a metric to quantify this configuration relative to the initiating SSUs and showed that SSU poising is highly enriched in mRNAs found in the synaptic compartment, suggesting regulation of the transition from poised to initiating SSUs in a subset of transcripts. In the mouse brain, SSU poising could be necessary for transcripts with slow elongation or start codon recognition rates and could allow fast and abundant translation in response to stimuli, such as during neuronal activity-induced bursts of protein synthesis. Supporting the latter hypothesis, we find that genes with high SSU poising ratios show higher translation efficiency as compared to those with lower SSU poising. One of the interesting gene candidates is *Camk2*α, which is known to undergo rapid dendritic translation upon synaptic stimulation and regulate excitatory synapse structure and function (Håvik et al. 2003; Miller et al. 2002; Ostroff et al. 2002; Chirillo et al. 2019; Yasuda, Hayashi, and Hell 2022). Thus, the SSU poised ratio could potentially reveal transcripts that are regulated at the stage of start codon recognition.

As discussed earlier (Archer et al. 2016), factors influencing and causing poising of SSUs may be multifaceted (e.g. mRNA sequence and structure, presence of IRES, SSU pausing). We found that the presence of uORFs is associated with less SSU poising, potentially by disassociating the SSUs after translation of the uORF. Similarly, the presence of uORFs can have an inhibitory effect on the translation of the downstream CDS. In line with previous reports (Calvo, Pagliarini, and Mootha 2009; Chew, Pauli, and Schier 2016; Johnstone, Bazzini, and Giraldez 2016), we observed globally a significant decrease in translational efficiency for transcripts harbouring a translated uORF.

### Cell-type and compartment specific translation regulation

When characterizing previously published lists of neuronal and glial enriched transcripts (Glock et al. 2021), we observed that neuronal transcripts have higher coverage of SSU and 80S than glial transcripts, potentially indicating abundance of neuronal over glial cells in the DG tissue, as shown before (Rieskamp et al. 2022). However, relative to RNA, neuronal genes showed higher translation and scanning efficiencies than those belonging to the glial category, metrics that are independent of cell-type abundance. Focusing on transcripts that are predominantly monosomal or polysomal provided further mechanistic details on their translation regulation. Neuronal transcripts preferring polysome translation showed higher SSU and 80S occupancy, whereas monosome-preferring transcripts showed lower SSU and 80S reads, implying reduced recruitment of PIC to these transcripts in order to maintain a low initiation and elongation rate. Indeed, monosome-preferring transcripts showed lower initiation rates along with lower translation efficiencies, as compared to polysome-preferring transcripts suggesting that monosome-preferring transcripts are more sparsely or selectively translated through a mechanism reducing PIC recruitment. The findings underscore the importance of transcriptome-wide mapping of both initiating and elongating ribosomes in providing an understanding of the plausible translation control mechanism of a specific pool of mRNAs.

## Conclusion

Here, we adapted and optimised RCP-seq for brain tissue, allowing the capture of SSUs, and analysis of translation initiation dynamics in mouse dentate gyrus and cerebral cortex. In tandem with total RNA and 80S analysis, we uncover cell type, and transcript-specific regulation at the scanning and elongation stages. We discover thousands of active uORFs associated with repressive function in the translation of the CDS. Additionally, we characterize a phenomenon of poised SSUs upstream of the translation initiation site of synaptically enriched RNAs, identifying it as a possible regulatory step during translation, underlying synapse maintenance and plasticity.

We anticipate that studies on scanning subunit occupancy will help to resolve long-standing questions on the role of initiation factors in the scanning process and how these influence the regulation of individual genes in mammalian brain function and dysfunction.

## Materials and Methods

### Animals

12 to 14-week-old C57BL/6 wild-type male mice were used. Mice were bred and housed in their home cages. Room temperature (22◦C (±1◦C)) and relative humidity (46±5%) were maintained. Mice had free access to water and food and were maintained on a 12-hour light/dark cycle. This research is approved by the Norwegian National Research Ethics Committee in compliance with EU Directive 2010/63/EU, ARRIVE guidelines.

### Crosslinking of HEK293T cells and ribosome complex profiling

HEK293T cells are cultured and maintained in DMEM supplemented with 10% FBS, penicillin and streptomycin and L-glutamine. Cells are grown in Ø 15 cm dishes up to 60-70% confluency prior to fixation. ∼3h before lysis the volume of media is made up to 20 mL.

For *in vitro* UV crosslinking, cells are UV cross-linked at 254 nm with 400 mJ/cm2 three times in a CL-1000 ultraviolet crosslinker (UVP). The plate is swirled between each round of irradiation for uniform crosslinking. Cells are then washed with 25 mL of ice-cold 1X PBS. The cells are scraped in a 1000 μL of ice-cold lysis buffer (10mM HEPES-KOH pH 7.5, 62.5 mM KCl, 2.5 mM MgCl_2_, 1 mM DTT, 1% Triton X 100, and protease inhibitor cocktail). The lysate is incubated in ice for 10 min and spun at 14000 RPM for 5 min to obtain the supernatant.

For formaldehyde crosslinking, 600 μL of 10% formaldehyde (0.3% final concentration) is added by tilting the dish to the pool of media and turning back the dishes to an even level. The cells are incubated on ice for 5 min with intermittent swirling. Formaldehyde is quenched by adding 600 μL of chilled 2.5 M glycine to media and incubated for 5 min, gently swirling after every minute. Cells are then washed with 25 mL of ice-cold 1X PBS. The cells are scraped in a 1000 μL of ice-cold lysis buffer. The lysate is incubated in ice for 10 min and spun at 14000 RPM for 5 min to obtain the supernatant.

For UV crosslinking after lysis, cells were washed with 20 mL ice-cold 1X PBS and scraped in a 1000 μL lysis buffer. The lysate was incubated in ice for 10 min and spun at 14000 RPM for 5 min. The supernatant in a 60mm dish is then UV cross-linked at 254 nm with 400 mJ/cm^2^ three times in a CL-1000 ultraviolet crosslinker (UVP). The dish was swirled between each round of irradiation for uniform crosslinking.

For RCP-sequencing, 10 OD lysates are digested with 70U of RNase1 for 45 min at 24C. The digestion is stopped with 28U of Superase In. The lysates are layered on 15-45% sucrose gradients and spun at 39000 RPM for 4 hours at 4 degree Celsius. Twenty fractions are collected of which RNA is extracted from fractions corresponding to 40S peak. De-crosslinking buffer (1% SDS, 10 mM EDTA, 10 mM Tris-HCl (pH 7.4), 10 mM glycine) is added to the 40S fractions. An equal volume of Phenol-Chloroform (pH 4.5) is added to the above mix and samples are incubated at 65 degree Celsius for 45 min at 1300 RPM shaking. The final RNA pellet is air-dried and resuspended in 12 μL of nuclease free water. RNA is processed according to the protocol mentioned later in the section.

### UV fixation of brain tissue and polysome profiling

Animals were anesthetized with urethane (1.5g/kg) and then sacrificed immediately. The whole brain was removed and positioned on a filter paper placed on a glass plate cooled with ice. After isolating both hippocampi, blood vessels and connective tissue were removed. The Cornu Ammonis (CA) region was separated from the dentate gyrus (DG) before placing the DG in microtubes in dry ice. After isolating both cerebral cortices, the white matter was carefully removed before placing the entire cortices in microtubes in dry ice. The brain samples were given a liquid nitrogen bath and then placed at -80 degrees Celsius for storage. It took a maximum of 10 minutes from sacrifice to sample storage to preserve RNA integrity. Long-term storage of tissue at -80 degrees Celsius was avoided to prevent RNA degradation.

Brain tissue was washed twice with ice-cold 1X PBS containing cycloheximide (0.1 mg/mL) before proceeding with the fixation/homogenisation. For formaldehyde fixation, whole tissues were incubated with 1mL formaldehyde solution for 5 min in ice. To neutralise formaldehyde, 1 M glycine was added and incubated for another 5 min. The tissue was then washed with 1X PBS twice before homogenisation. For UV crosslinking, the tissue was first lysed and the lysate was exposed to UV. Tissue homogenisation was done in lysis buffer (50 mM Tris-Cl, pH 7.4; 100 mM KCl; 5 mM MgCl_2_; EDTA-free protease inhibitor cocktail; 1 mM DTT; 0.1 mg/mL cycloheximide, 40 U/mL Superase Inhibitor) in a dounce homogenizer with 15 strokes of loose pistol and 15 strokes of the tight piston on ice. The volume of the lysis buffer used depended on the size of the tissue. For the cortex (one hemisphere), 0.5 mL was used. For DG regions, 0.3 mL was used. For one replicate, one cortical hemisphere (∼17ug RNA) was used and 10 DGs (∼9ug RNA) were used. The homogenate was transferred to an RNase-free 1.5 mL tube and 1% NP40 was added to the homogenate, mixed well, and incubated in ice for 10 min. The homogenate was spun at 2000 g for 10 min at 4 degrees Celsius. The clear supernatant (S1) was collected and then spun at 20000 g for 10 min at 4 degrees Celsius. The clear supernatant (S2) obtained was transferred to a 3.5 cm tissue culture dish placed on a bed of ice slush. The supernatant was UV cross-linked at 254 nm with 400 mJ/cm^2^ three times in a CL-1000 ultraviolet crosslinker (UVP). The plate was swirled between each round of irradiation for uniform crosslinking. The RNA concentration was estimated by the absorbance using both qubit and nanodrop. This can be done before or after UV irradiation. The lysate was layered over a 10-50% sucrose gradient (10 mM Tris-Cl pH7.4, 100 mM KCl, 5 mM MgCl_2_, 2 mM DTT) and spun at 40000 RPM for 2-4 hours at 4 degrees Celsius in an ultracentrifuge. The gradients were then run through a UV detector-fractionator (Biocomp).

### Ribosome complex profiling: library prep and sequencing

RNase digestion was done as previously described (Liu et al. 2019). 5U of RNase 1 was used for one unit of absorbance at A260 for a total volume of 300 μL brain lysate. RNase digestion for UV crosslinked lysates was done at 25 degrees Celsius for 45 min. The digestion was stopped by the addition of 50U of Superase Inhibitor. For total RNA sequencing, 50-100 μL of lysate was kept aside before the digestion step. 10-50% sucrose gradients were made 45 min before the ultracentrifugation in polycarbonate tubes (Science Services, Germany, S7030). 450-500 μL of digested lysates were then layered over the gradients and run in SW41 Ti rotor at 40000 rpm for 3 hours. The gradients were then run through a UV detector-fractionator (Biocomp) and 20 fractions were collected. Fractions corresponding to the 40S and 80S peaks were used for RNA extraction and library preparation.

RNA extraction was done using TRIzol LS and in the final step, the RNA pellet was resuspended in 10 μl of nuclease-free water. The RNA sample was then run on a 15% polyacrylamide gel for size selection for the 40S and 80S footprints using RNA denaturing dye, with an ultra-low DNA ladder as the marker (Invitrogen: 10597012). For 40S footprints, gel from 20-80 nt was cut and for 80S footprints, gel from 20-50 nt was cut. The gel pieces were crushed and resuspended in 500 μL of 0.3M Sodium acetate solution overnight, at 4 degrees Celsius, and then filtered through 0.4 μm cellulose filters. RNA was precipitated overnight at -20 degrees Celsius in the presence of 2 μl of glycoblue (Life technologies: AM9516) and 375 μL isopropanol and resuspended in 10 μl of nuclease-free water. The size selected footprints were 3’ dephosphorylated using PNK for 2 hours at 37 degrees Celsius in a final volume of 20 μl. After dephosphorylation, the RNA was concentrated and cleaned by the Oligo clean and concentrator kit (Zymo Research) in a final volume of 14 μL in nuclease-free water. The RNA footprints were then rRNA depleted with oligos from siTOOLS (riboPOOL riboseq h/m/r) and eluted in a final volume of 10 μl. Small RNA libraries were made using the Takara SMARTer smRNA kit and the manufacturer’s protocol was followed. PCR cycles were based on the starting concentration of RNA. Total RNA libraries were prepared using the Takara SMARTer total RNA HI mammalian kit and the manufacturer’s protocol was followed. Libraries containing primers or primer dimers were cleaned up using RNA XP clean-up beads (catalogue #A63987). The size and molarity of the libraries were estimated by obtaining bioanalyzer profiles (Agilent). Libraries were run as 100 bp single end on the NOVAseq 6000 platform up to a depth of 100M reads for each small RNA library or 30M reads for the total RNA library.

### Ribosome complex profiling: analysis

The repository at https://git.app.uib.no/valenlab/brain_dg_cortex_rcp and https://git.app.uib.no/valenlab/hek293_preeti contains all the code that was used for analysis and all of the processed data and figures are available there for inspection. All figures and processed tables are present in the repository. For the data processing, we used ORFik (1.19.3) as shown in the script “0_preprocess.R”. Analysis uses the latest at the time, gencode mouse release M31 (GRCm39 v107). ORFik pipeline uses fastp software for trimming and STAR for alignment. Paired-end fastq reads are initially aligned to the contaminants - phix, rRNA, ncRNA, tRNA, and finally to the genome. Important options that were set are adapter.sequence = "AAAAAAAAAA", trim.front = 3, min.length = 20. Aligned data is processed using ORFik and custom scripts available in the data repository. We restricted genes with multiple transcripts to a single transcript by selecting the one with the longest coding sequence. For statistical testing, we used cutoffs of log2 fold change of 1 or -1 and 0.05 as the significance level. Upstream ORFs were detected using the ORFik function “findUORFs” which by default searches for HTG start codons (not GAG), but we used only ATG as start codons. Furthermore, we filtered out these uORFs that don’t have any values over SSU/80S/RNA. We restricted further to one uORF per transcript by selecting the one with the highest translational efficiency.

For comparisons at the global-level, we calculate measures produced per library using FPKM values instead of raw counts and average replicates, furthermore, we normalise using reads per million reads for that particular group of genes to gain a comprehensive perspective. As we can see below, uORF measures are adapted from gene measures by changing the perspective of CDS to that of the uORF, and 5’ leader and 3’ trailer are becoming regions upstream/downstream of uORF CDS, but within boundaries of the transcript mRNA.

### Formulae used in this study

*Scanning efficiency (SE) = SSU on leaders / RNA on mRNA (leader + CDS + trailer)*

*Translation efficiency (TE) = 80S on CDS / RNA on mRNA*

*Initiation rate (IR) = 80S on CDS / SSU on leaders*

*uORF TE = 80S on uORF/ RNA on uORFs*

*uORF SSU consumption rate = SSU upstream of uORF / SSU downstream of uORF*

*uORF ratio = 80S of uORF / 80S of CDS*

*SSU poising: Long SSU (60-65nt) at -46: -36 relative to TIS / Short SSU (25-35nt) at -14: -12 relative to TIS*

*Poised SSUs/RNA: Long SSU (60-65nt) at -46: -36 relative to TIS / RNA on mRNA*

### Statistics and reproducibility

Animal experiments were done in three independent replicates for each tissue (n = 3). Only one replicate was performed for HEK293T experiments (n = 1). For boxplot figures t-test statistics were used to compare means of the distributions with corrected p-values using Benjamini-Hochberg method.

## Data availability

Raw reads are published under accession number PRJEB72224 in the European Nucleotide Archive.

## Author contributions

PMK, FPP, EV and CRB designed the experiments. FPP performed the mice brain tissue dissection and PMK performed the ribosome complex profiling experiments. KL and EV conducted the bioinformatic analysis. PMK, KL and EV interpreted the data. All authors discussed the results and contributed to writing the manuscript.

## Acknowledgements

FPP was funded by the L.Meltzers Høyskolefond 2022 - Ekstratildeling - for the project “Molecular control of protein synthesis underlying the plasticity of neural communication in the mammalian brain” (Project #103517122). Research in the CRB lab is supported by the Trond Mohn Research Foundation (TMS2021TMT04). PK and EV are funded by the Norwegian Cancer Society (Project #190290). The Genomics Core Facility (GCF) at the University of Bergen, which is a part of the NorSeq consortium, provided services on sequencing of small and total RNA libraries on NOVAseq 6000 platform; GCF is supported in part by major grants from the Research Council of Norway (grant no. 245979/F50) and Bergen Research Foundation (BFS) (grant no. BFS2017TMT04 and BFS2017TMT08)”.

## Supplementary figures and tables

**Supplementary Figure 1:**
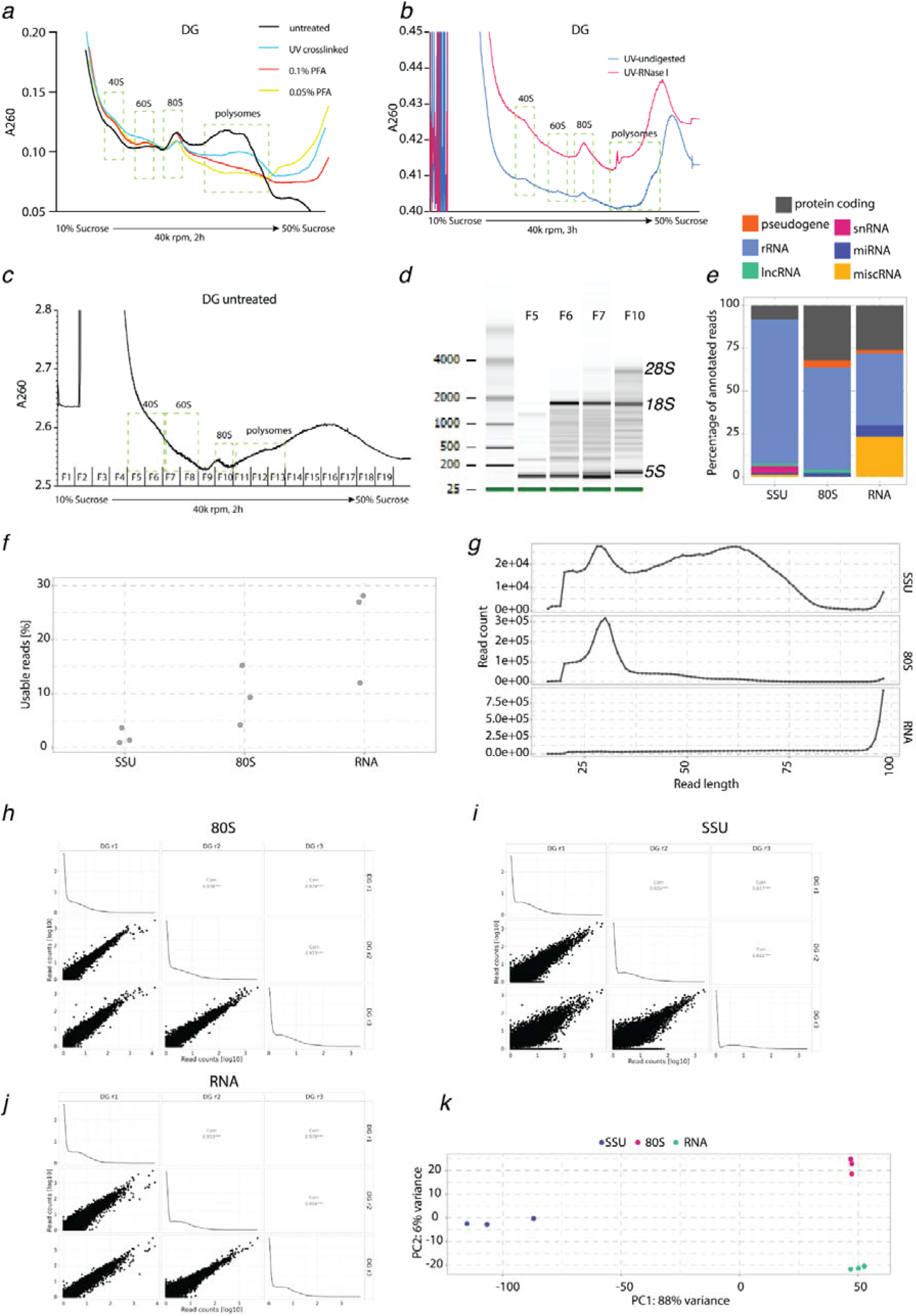
Quality assessment of the UV-crosslinking method for RCP-seq (Related to Figure 1) **a**: Amount of crosslinking for polysomes for three different cross-linking methods: formaldehyde, UV or non-crosslinked DG samples. **b**: Polysome profiles of UV-crosslinked, with or without RNase I digestion to denote increase in 80S for digested samples for DG **c**: A UV trace of DG polysomes treated only with cycloheximide (no crosslinking) from a 10-15% sucrose gradient. 20 fractions were collected. Ribosomal subunits, 80S and polysomes are indicated. **d**: Bioanalyzer image of total RNA isolated from fractions from the above profile. From this image, fractions 5 and 6 contain 40S. **e**: Percent content displaying the mapping of SSU, 80S and total RNA counts to the type of transcript. **f**: Percent reads that are used for the gene expression analysis for SSU, 80S and RNA libraries. **g**: Footprint length distribution for the DG for SSU, 80S and RNA libraries. **h**: Pearson’s correlation for 80S among replicates in DG **i**: Pearson’s correlation for SSU among replicates in DG **j**: Pearson’s correlation for RNA among replicates in DG **k**: PCA plot for SSU, 80S and RNA libraries from DG

**Supplementary Figure 2:**
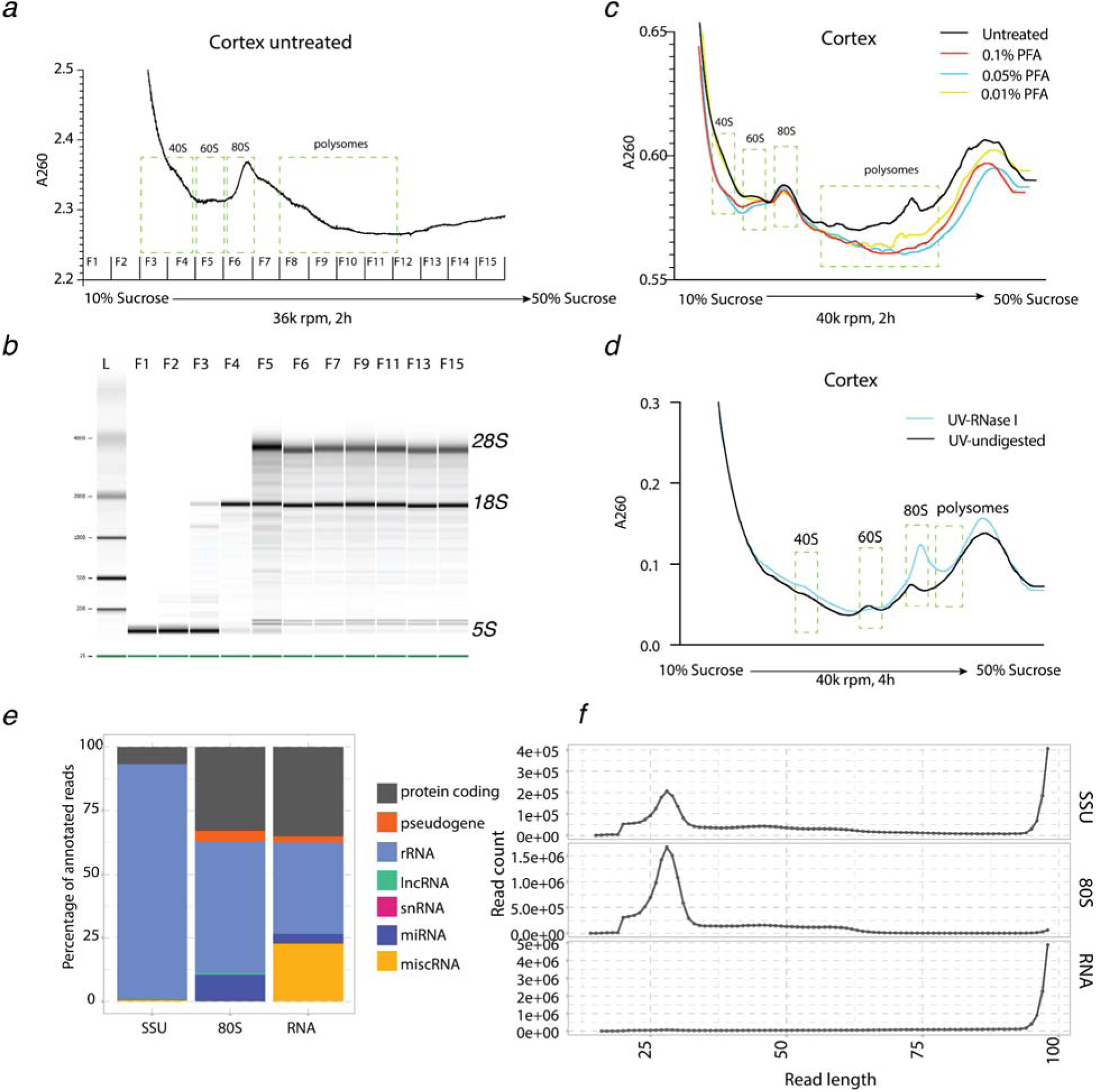
UV-crosslinking method for RCP-seq in the cortical tissue (related to figure 2) **a**: A UV trace of cortical polysomes treated only with cycloheximide (no crosslinking) from a 10-50% sucrose gradient. 20 fractions were collected out of which 15 fractions are shown. **b**: Bioanalyzer image of total RNA isolated from fractions from the above profile. From this image, fractions 3 and 4 may contain 40S. **c**: Amount of crosslinking for polysomes for three different paraformaldehyde concentrations vs non-crosslinked. **d:** Polysome profiles of UV-crosslinked, with or without RNase I digestion to denote an increase in the 80S for digested samples for cortical tissue **e**: Percent content displaying the mapping of SSU, 80S and total RNA counts to the type of transcript. **f:** Footprint length distribution for the DG for SSU, 80S and RNA libraries.

**Supplementary Figure 3:**
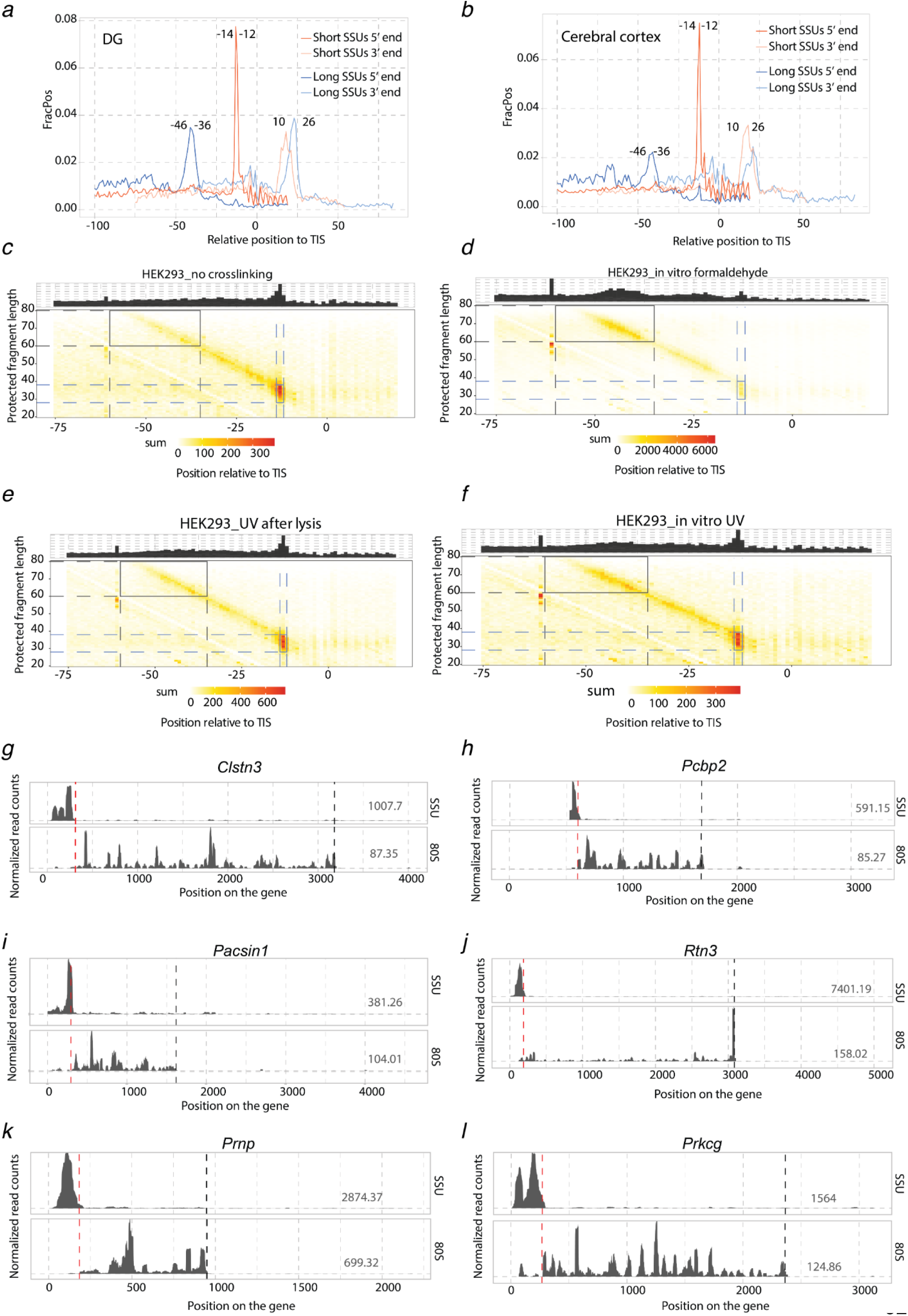
Estimation of SSU poising in 5’ leaders of transcripts from the DG and th cerebral cortex (Related to Figure 3) **a** and **b:** Frequency distribution of SSU footprints based on their 5’ end position on the leaders for the DG (**a**) and for the cerebral cortex (**b**). Numbers on the plots indicate nucleotide positions on the leaders relative to the TIS/start codon. **c**-**f:** Footprint length distribution for the 5’ end of SSU fragments in HEK293T cells relative to the TIS, highlighting initiatin SSUs (at -12nt) and poised SSUs (−36: -60nt) for different crosslinking condition, for non-crosslinked (**c**), for formaldehyde crosslinked (**d**), for UV crosslinked after lysis (**e**), for in vitro UV crosslinking (**f**) **g-l**: Single gene profile from DG for SSU and 80S, without intronic regions, and plotted for the longest isoform of *Clstn* (Calsyntenin3, **g**), *Pcbp2* (Poly (rC) binding protein 2, **h**), *Pacsin1* (Protein Kinase C And Casein Kinase Substrate In Neurons 1, **i**), *Rtn3* (Reticulon 3, **j**), *Prnp* (Prion protein, **k**), *Prkcg* (Protein kinase C, gamma, **l**). Numbers in the plot indicate FPKM values. Dotted lines indicate TIS (red) and TTS (black).

**Supplementary figure 4:**
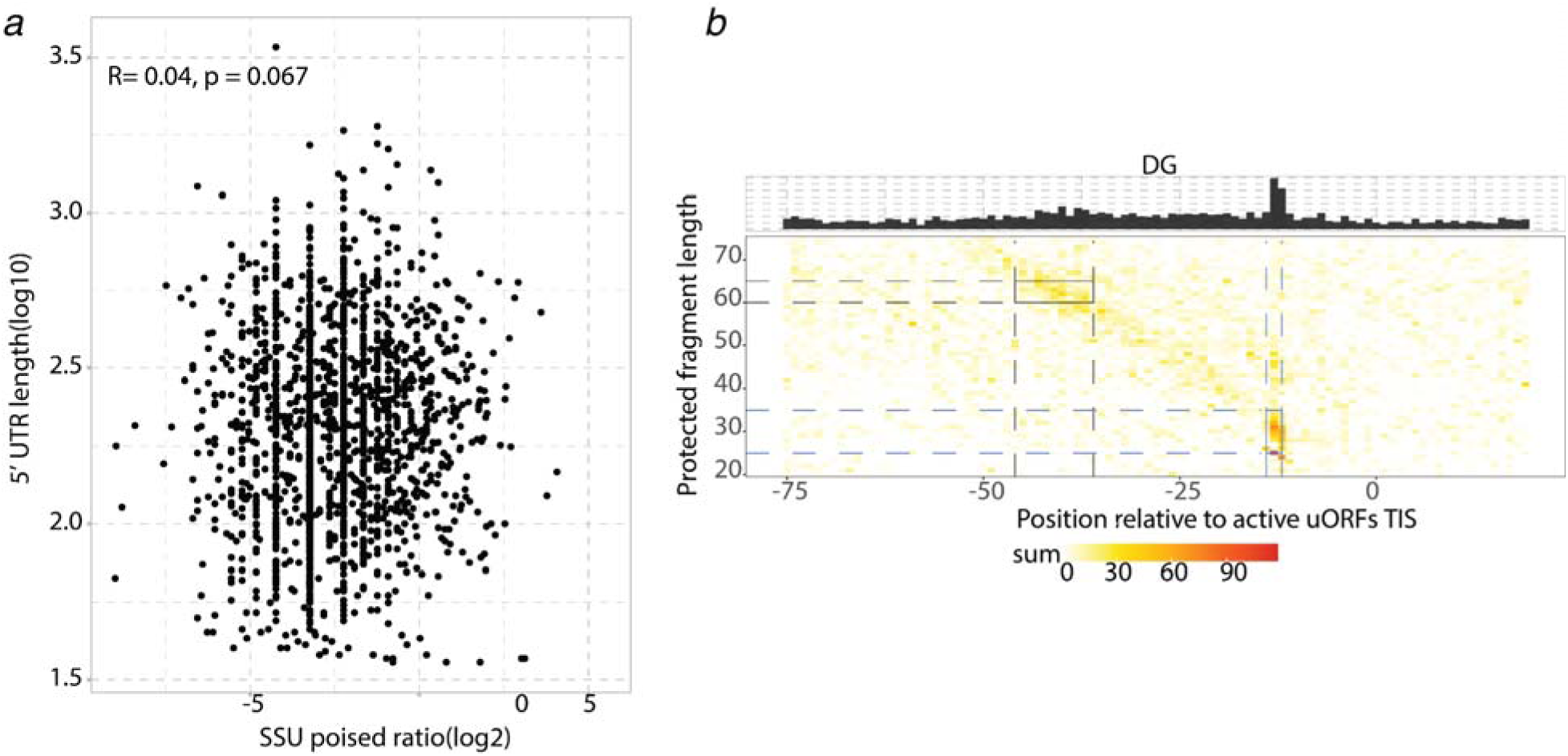
Effect of leaders on SSU poising (related to figure 4) **a**: Log2 of SSU poised ratio *vs*. the length of the 5’ UTRs for transcripts from DG. **b**: Footprint length distribution for the 5’ end SSU fragments relative to TIS of active uORFs in the DG tissue, highlightin initiating SSUs (at -12nt) and poised SSUs (−46: -42nt).

**Supplementary Table 1:** Sequencing depth, contaminants mapping.

**Supplementary Table 2:** overview of descriptions of queued SSU.

| Reference | Model system | Localization | Suggested mechanism? | Detection method |
| --- | --- | --- | --- | --- |
| (Ivanov et al. 2018) | HEK293T cells, reporter assays | SSU queuing on uORF, upstream of paused ribosome | high polyamines, elongation pausing (over PPW motif) during translation of the uORF | Model |
| (Kearse et al. 2019) | HeLa cells, reporter assays | SSU queuing on ORF, upstream of paused ribosome | non-AUG start codons, slowed elongating ribosomes | Model |
| (Archer et al. 2016) | Yeast | SSU queuing on leaders with 5' end -30nt rel to TIS | local structural or sequence features of mRNAs | TCP-seq |
| (Wagner et al. 2020) | Yeast and HEK293T | In yeast, SSU queuing on leaders -30nt rel to TIS, for HEK293T SSU queuing on leaders -60 to -30nt rel to TIS |  | Sel-TCP seq and TCP-seq |
| (Bohlen et al. 2020) | NIH 3T3 | SSU queuing on leaders -120nt to -60nt rel to TIS | 80S ribosomes stalled at >12 codons downstream of TIS | Harringtonine treatment and TCP-seq |

**Supplementary Table 3:** List of SSU poised-up genes in both DG and cortex.

**Supplementary Table 4:** Active uORFs discovered in the study in both DG and cortex.

## References

Amorim, Inês S., Sonal Kedia, Stella Kouloulia, Konstanze Simbriger, Ilse Gantois, Seyed Mehdi Jafarnejad, Yupeng Li, et al. 2018. “Loss of eIF4E Phosphorylation Engenders Depression-like Behaviors via Selective mRNA Translation.” The Journal of Neuroscience: The Official Journal of the Society for Neuroscience 38 (8): 2118–33.

Archer, Stuart K., Nikolay E. Shirokikh, Traude H. Beilharz, and Thomas Preiss. 2016. “Dynamics of Ribosome Scanning and Recycling Revealed by Translation Complex Profiling.” Nature 535 (7613): 570–74.

Biever, Anne, Caspar Glock, Georgi Tushev, Elena Ciirdaeva, Tamas Dalmay, Julian D. Langer, and Erin M. Schuman. 2020. “Monosomes Actively Translate Synaptic mRNAs in Neuronal Processes.” Science 367 (6477). 10.1126/science.aay4991.

Bohlen, Jonathan, Kai Fenzl, Günter Kramer, Bernd Bukau, and Aurelio A. Teleman. 2020. “Selective 40S Footprinting Reveals Cap-Tethered Ribosome Scanning in Human Cells.” Molecular Cell 79 (4): 561–74.e5.

Bramham, Clive R., and David G. Wells. 2007. “Dendritic mRNA: Transport, Translation and Function.” Nature Reviews. Neuroscience 8 (10): 776–89.

Brito Querido, Jailson, Masaaki Sokabe, Irene Díaz-López, Yuliya Gordiyenko, Christopher S. Fraser, and V. Ramakrishnan. 2024. “The Structure of a Human Translation Initiation Complex Reveals Two Independent Roles for the Helicase eIF4A.” Nature Structural & Molecular Biology 31 (3): 455–64.

Calvo, Sarah E., David J. Pagliarini, and Vamsi K. Mootha. 2009. “Upstream Open Reading Frames Cause Widespread Reduction of Protein Expression and Are Polymorphic among Humans.” Proceedings of the National Academy of Sciences of the United States of America 106 (18): 7507– 12.

Cao, Ruifeng, Christos G. Gkogkas, Nuria de Zavalia, Ian D. Blum, Akiko Yanagiya, Yoshinori Tsukumo, Haiyan Xu, et al. 2015. “Light-Regulated Translational Control of Circadian Behavior by eIF4E Phosphorylation.” Nature Neuroscience 18 (6): 855–62.

Chew, Guo-Liang, Andrea Pauli, and Alexander F. Schier. 2016. “Conservation of uORF Repressiveness and Sequence Features in Mouse, Human and Zebrafish.” Nature Communications 7 (May):11663.

Chirillo, Michael A., Mikayla S. Waters, Laurence F. Lindsey, Jennifer N. Bourne, and Kristen M. Harris. 2019. “Local Resources of Polyribosomes and SER Promote Synapse Enlargement and Spine Clustering after Long-Term Potentiation in Adult Rat Hippocampus.” Scientific Reports 9 (1): 3861.

Darnell, Jennifer C., Sarah J. Van Driesche, Chaolin Zhang, Ka Ying Sharon Hung, Aldo Mele, Claire E. Fraser, Elizabeth F. Stone, et al. 2011. “FMRP Stalls Ribosomal Translocation on mRNAs Linked to Synaptic Function and Autism.” Cell 146 (2): 247–61.

Dever, Thomas E., Ivaylo P. Ivanov, and Alan G. Hinnebusch. 2023. “Translational Regulation by uORFs and Start Codon Selection Stringency.” Genes & Development 37 (11-12): 474–89.

Giess, Adam, Yamila N. Torres Cleuren, Håkon Tjeldnes, Maximilian Krause, Teshome Tilahun Bizuayehu, Senna Hiensch, Aniekan Okon, Carston R. Wagner, and Eivind Valen. 2020. “Profiling of Small Ribosomal Subunits Reveals Modes and Regulation of Translation Initiation.” Cell Reports 31 (3): 107534.

Glock, Caspar, Anne Biever, Georgi Tushev, Belquis Nassim-Assir, Allison Kao, Ina Bartnik, Susanne Tom Dieck, and Erin M. Schuman. 2021. “The Translatome of Neuronal Cell Bodies, Dendrites, and Axons.” Proceedings of the National Academy of Sciences of the United States of America 118 (43). 10.1073/pnas.2113929118.

Hafner, Markus, Maria Katsantoni, Tino Köster, James Marks, Joyita Mukherjee, Dorothee Staiger, Jernej Ule, and Mihaela Zavolan. 2021. “CLIP and Complementary Methods.” Nature Reviews Methods Primers 1 (1): 1–23.

Håvik, Bjarte, Håvard Røkke, Kjetil Bårdsen, Svend Davanger, and Clive R. Bramham. 2003. “Bursts of High-Frequency Stimulation Trigger Rapid Delivery of Pre-Existing Alpha-CaMKII mRNA to Synapses: A Mechanism in Dendritic Protein Synthesis during Long-Term Potentiation in Adult Awake Rats.” The European Journal of Neuroscience 17 (12): 2679–89.

Hien, Annie, Gemma Molinaro, Botao Liu, Kimberly M. Huber, and Joel D. Richter. 2020. “Ribosome Profiling in Mouse Hippocampus: Plasticity-Induced Regulation and Bidirectional Control by TSC2 and FMRP.” Molecular Autism 11 (1): 78.

Hinnebusch, Alan G. 2014. “The Scanning Mechanism of Eukaryotic Translation Initiation.” Annual Review of Biochemistry 83 (January):779–812.

Holt, Christine E., Kelsey C. Martin, and Erin M. Schuman. 2019. “Local Translation in Neurons: Visualization and Function.” Nature Structural & Molecular Biology 26 (7): 557–66.

Ingolia, Nicholas T., Sina Ghaemmaghami, John R. S. Newman, and Jonathan S. Weissman. 2009. “Genome-Wide Analysis in Vivo of Translation with Nucleotide Resolution Using Ribosome Profiling.” Science 324 (5924): 218–23.

Iordanov, Mihail S., Remy J. Choi, Olga P. Ryabinina, Thanh-Hoai Dinh, Robert K. Bright, and Bruce E. Magun. 2002. “The UV (Ribotoxic) Stress Response of Human Keratinocytes Involves the Unexpected Uncoupling of the Ras-Extracellular Signal-Regulated Kinase Signaling Cascade from the Activated Epidermal Growth Factor Receptor.” Molecular and Cellular Biology 22 (15): 5380– 94.

Iordanov, M. S., D. Pribnow, J. L. Magun, T. H. Dinh, J. A. Pearson, and B. E. Magun. 1998. “Ultraviolet Radiation Triggers the Ribotoxic Stress Response in Mammalian Cells.” The Journal of Biological Chemistry 273 (25): 15794–803.

Ivanov, Ivaylo P., Byung-Sik Shin, Gary Loughran, Ioanna Tzani, Sara K. Young-Baird, Chune Cao, John F. Atkins, and Thomas E. Dever. 2018. “Polyamine Control of Translation Elongation Regulates Start Site Selection on Antizyme Inhibitor mRNA via Ribosome Queuing.” Molecular Cell 70 (2): 254–64.e6.

Johnstone, Timothy G., Ariel A. Bazzini, and Antonio J. Giraldez. 2016. “Upstream ORFs Are Prevalent Translational Repressors in Vertebrates.” The EMBO Journal 35 (7): 706–23.

Kearse, Michael G., Daniel H. Goldman, Jiou Choi, Chike Nwaezeapu, Dongming Liang, Katelyn M. Green, Aaron C. Goldstrohm, Peter K. Todd, Rachel Green, and Jeremy E. Wilusz. 2019. “Ribosome Queuing Enables Non-AUG Translation to Be Resistant to Multiple Protein Synthesis Inhibitors.” Genes & Development 33 (13-14): 871–85.

Kute, Preeti Madhav, Omar Soukarieh, Håkon Tjeldnes, David-Alexandre Trégouët, and Eivind Valen. 2021. “Small Open Reading Frames, How to Find Them and Determine Their Function.” Frontiers in Genetics 12:796060.

Larsson, Ola, Bin Tian, and Nahum Sonenberg. 2013. “Toward a Genome-Wide Landscape of Translational Control.” Cold Spring Harbor Perspectives in Biology 5 (1): a012302.

Liu, Botao, Gemma Molinaro, Huan Shu, Emily E. Stackpole, Kimberly M. Huber, and Joel D. Richter. 2019. “Optimization of Ribosome Profiling Using Low-Input Brain Tissue from Fragile X Syndrome Model Mice.” Nucleic Acids Research 47 (5): e25.

Merrick, William C., and Graham D. Pavitt. 2018. “Protein Synthesis Initiation in Eukaryotic Cells.” Cold Spring Harbor Perspectives in Biology 10 (12). 10.1101/cshperspect.a033092.

Miller, Stephan, Masahiro Yasuda, Jennifer K. Coats, Ying Jones, Maryann E. Martone, and Mark Mayford. 2002. “Disruption of Dendritic Translation of CaMKIIα Impairs Stabilization of Synaptic Plasticity and Memory Consolidation.” Neuron 36 (3): 507–19.

Moon, Stephanie L., Nahum Sonenberg, and Roy Parker. 2018. “Neuronal Regulation of eIF2α Function in Health and Neurological Disorders.” Trends in Molecular Medicine 24 (6): 575–89.

Oliveira, Mauricio M., and Eric Klann. 2022. “eIF2-Dependent Translation Initiation: Memory Consolidation and Disruption in Alzheimer’s Disease.” Seminars in Cell & Developmental Biology 125 (May):101–9.

Ostroff, Linnaea E., John C. Fiala, Brenda Allwardt, and Kristen M. Harris. 2002. “Polyribosomes Redistribute from Dendritic Shafts into Spines with Enlarged Synapses during LTP in Developing Rat Hippocampal Slices.” Neuron 35 (3): 535–45.

Patil, Sudarshan, Kleanthi Chalkiadaki, Tadiwos F. Mergiya, Konstanze Krimbacher, Inês S. Amorim, Shreeram Akerkar, Christos G. Gkogkas, and Clive R. Bramham. 2023. “eIF4E Phosphorylation Recruits β-Catenin to mRNA Cap and Promotes Wnt Pathway Translation in Dentate Gyrus LTP Maintenance.” iScience. https://www.cell.com/iscience/pdf/S2589-0042(23)00726-5.pdf.

Rieskamp, Joshua D., Patricia Sarchet, Bryon M. Smith, and Elizabeth D. Kirby. 2022. “Estimation of the Density of Neural, Glial, and Endothelial Lineage Cells in the Adult Mouse Dentate Gyrus.” Neural Regeneration Research 17 (6): 1286–92.

Ruiz-Orera, Jorge, and M. Mar Albà. 2019. “Conserved Regions in Long Non-Coding RNAs Contain Abundant Translation and Protein-RNA Interaction Signatures.” NAR Genomics and Bioinformatics 1 (1): e2.

Sharma, Vijendra, Rapita Sood, Abdessattar Khlaifia, Mohammad Javad Eslamizade, Tzu-Yu Hung, Danning Lou, Azam Asgarihafshejani, et al. 2020. “eIF2α Controls Memory Consolidation via Excitatory and Somatostatin Neurons.” Nature 586 (7829): 412–16.

Shirokikh, Nikolay E., Yulia S. Dutikova, Maria A. Staroverova, Ross D. Hannan, and Thomas Preiss. 2019. “Migration of Small Ribosomal Subunits on the 5’ Untranslated Regions of Capped Messenger RNA.” International Journal of Molecular Sciences 20 (18): 4464.

Sokabe, Masaaki, and Christopher S. Fraser. 2019. “Toward a Kinetic Understanding of Eukaryotic Translation.” Cold Spring Harbor Perspectives in Biology 11 (2). 10.1101/cshperspect.a032706.

Sun, Chao, and Erin Schuman. 2023. “A Multi-Omics View of Neuronal Subcellular Protein Synthesis.” Current Opinion in Neurobiology 80 (June):102705.

Thomson, Sophie R., Sang S. Seo, Stephanie A. Barnes, Susana R. Louros, Melania Muscas, Owen Dando, Caoimhe Kirby, et al. 2017. “Cell-Type-Specific Translation Profiling Reveals a Novel Strategy for Treating Fragile X Syndrome.” Neuron 95 (3): 550–63.e5.

Wagner, Susan, Jonathan Bohlen, Anna Herrmannova, Jan Jelínek, Thomas Preiss, Leoš Shivaya Valášek, and Aurelio A. Teleman. 2022. “Selective Footprinting of 40S and 80S Ribosome Subpopulations (Sel-TCP-Seq) to Study Translation and Its Control.” Nature Protocols 17 (10): 2139–87.

Wagner, Susan, Anna Herrmannová, Vladislava Hronová, Stanislava Gunišová, Neelam D. Sen, Ross D. Hannan, Alan G. Hinnebusch, Nikolay E. Shirokikh, Thomas Preiss, and Leoš Shivaya Valášek. 2020. “Selective Translation Complex Profiling Reveals Staged Initiation and Co-Translational Assembly of Initiation Factor Complexes.” Molecular Cell 79 (4): 546–60.e7.

Yasuda, Ryohei, Yasunori Hayashi, and Johannes W. Hell. 2022. “CaMKII: A Central Molecular Organizer of Synaptic Plasticity, Learning and Memory.” Nature Reviews. Neuroscience 23 (11): 666–82.

